# Executive and language systems competitively shape predictive processing during reading

**DOI:** 10.64898/2026.10.23.761476

**Authors:** Sandra Martin, Anna-Lisa Schuler, Ole Numssen, Niklas Kroner-Weigl, Merle M. Schuckart, Jonas Obleser, Gesa Hartwigsen

## Abstract

When we read, our brains constantly generate predictions about upcoming words, which is thought to make comprehension faster and more efficient. Yet it remains unclear whether this predictive mechanism relies on the same domain-general executive resources the brain uses for other demanding cognitive tasks, or operates largely independently within the language system. Here we tested this question using transcranial magnetic stimulation (TMS) in twenty-five healthy adults, who performed self-paced naturalistic reading, an n-back task, and a dual task of concurrent reading and n-back, allowing us to place competing demands on cognitive resources. In three sessions, we stimulated the left angular gyrus alone, the angular gyrus together with dorsolateral prefrontal cortex, a hub of executive control, or delivered sham stimulation, and used computational modeling of decision-making to isolate how each brain state altered the underlying cognitive processes. Combined stimulation of the angular gyrus and prefrontal cortex selectively weakened the reading-time costs of unexpected words, but only when reading competed with the working-memory task, revealing that linguistic prediction draws on executive resources specifically under cognitive load. Computational models showed that the relationship was bidirectional: Unpredictable words also disrupted concurrent working-memory decisions, slowing the accumulation of evidence and increasing response caution. Together, these findings show that predictive language processing is not a fixed, encapsulated function but is dynamically coupled to executive control when cognitive demands compete. Our results provide evidence that prediction during language comprehension is embedded within a broader, demand-sensitive cognitive architecture that flexibly coordinates linguistic and executive resources.

## Introduction

Predictive processing is increasingly viewed as a unifying computational principle across perception and cognition, whereby the brain continuously generates forward models of expected input to optimize processing efficiency [1]. Efficient language comprehension is thought to rely on this cognitive ability to generate probabilistic predictions about upcoming input based on the already perceived content [2,3]. In reading, this means that comprehenders continuously use preceding linguistic context to anticipate upcoming words, with less expected words typically incurring greater processing cost, reflected in longer reading times [4,5]. This relationship is commonly formalized as lexical surprisal, the negative log-probability of a word conditioned on its context, which has been repeatedly shown to predict word-level reading times across both eye-tracking and self-paced reading paradigms [6,7]. Critically, recent evidence suggests that the magnitude of the surprisal effect is attenuated when concurrent (non-verbal) cognitive demands compete for executive resources [8–10]. Such competing demands are common in naturalistic reading, where comprehenders frequently read while concurrently monitoring for target information or updating working memory, yet the neural systems that give rise to this load-dependent modulation, and how they respond to direct perturbation, remain poorly understood [11,12].

Two candidate regions are of particular interest for this interaction. Neuroimaging research has linked the left angular gyrus (AG) to semantic and contextual integration during reading, consistent with a role in combining incoming lexical information with sentence- and discourse-level context [13–16]. Electrophysiological work also implicates the left AG in anticipating upcoming events, possibly through re-orienting attention toward relevant contextual information [17,18]. This suggests that generating expectations may also be part of the AG’s broader role in sensemaking.

The left dorsolateral prefrontal cortex (DLPFC), in contrast, is consistently implicated in the executive control of working memory, particularly in the updating and monitoring of task-relevant information under increasing cognitive load [19,20]. As part of large-scale domain-general executive networks, it supports the maintenance and manipulation of information across verbal and non-verbal task content, and its recruitment increases with the cognitive demand for active monitoring. During reading specifically, this monitoring function has been linked to verbal working memory [21,22]. Whether and how this domain-general monitoring system interacts with AG’s proposed role in building context-based expectations, however, remains unresolved, particularly when both are taxed simultaneously. One possibility is that processing prediction errors, such as those elicited by low-probability words, draw on executive resources, such that the AG’s contribution to prediction becomes increasingly dependent on DLPFC-mediated control as cognitive load rises.

Non-invasive brain stimulation (NIBS) provides a way to probe the functional relevance of these proposed roles under varying cognitive demands. However, most NIBS studies on reading have relied on single-word paradigms, such as phonological, lexical, or semantic judgment tasks, isolating specific linguistic sub-processes rather than natural reading itself [for reviews see 23,24]. One study that used a natural reading paradigm in combination with repetitive transcranial magnetic stimulation (rTMS) over left AG showed that disrupting its activity impairs contextual integration and memory for coherent narratives [25], while another study demonstrated that facilitatory transcranial direct current stimulation speeds semantic integration of related word pairs [26]. Moreover, a TMS study from the auditory domain has directly linked the left AG to word predictability, showing that its disruption reduces the comprehension benefit of predictable input during degraded speech [27]. Together, these findings suggest a key role of left AG in both contextual integration and predictability, motivating a direct test of its relevance for moment-to-moment prediction during natural reading.

Evidence for the contribution of the DLPFC to working memory is more extensive and consistent. Numerous studies in healthy adults have shown that rTMS to the left DLPFC reliably modulates working-memory performance, for instance during the n-back paradigm, with high-frequency protocols and higher task demands usually leading to improved performance [for reviews see 28,29]. Because reading itself is thought to draw substantially on verbal working-memory resources [21], DLPFC stimulation offers a way to test whether modulating domain-general executive control also alters performance on a task with linguistic content. However, no study to date has combined AG and DLPFC stimulation within the same design to test whether their respective contributions to prediction and executive control interact when cognitive load is manipulated directly, providing new insights into the mechanistic underpinnings of natural language comprehension.

The present study addresses this gap. Across three sessions, participants performed three tasks: self-paced reading of naturalistic texts alone, an n-back task, and a dual task of concurrent reading and n-back. In each session, they either received 1 Hz rTMS to left AG before the experiment, combined 1 Hz AG rTMS before and 10 Hz rTMS to left DLPFC during the experiment, or sham stimulation (see Fig. 1). This design allowed us to test whether modulating AG and DLPFC function induces effects on reading and working memory that depend on concurrent cognitive load. Moreover, the dual task provided a way to tax both language processing and working memory systems simultaneously and to test whether their relative contributions shift with cognitive load.

**Figure 1.**
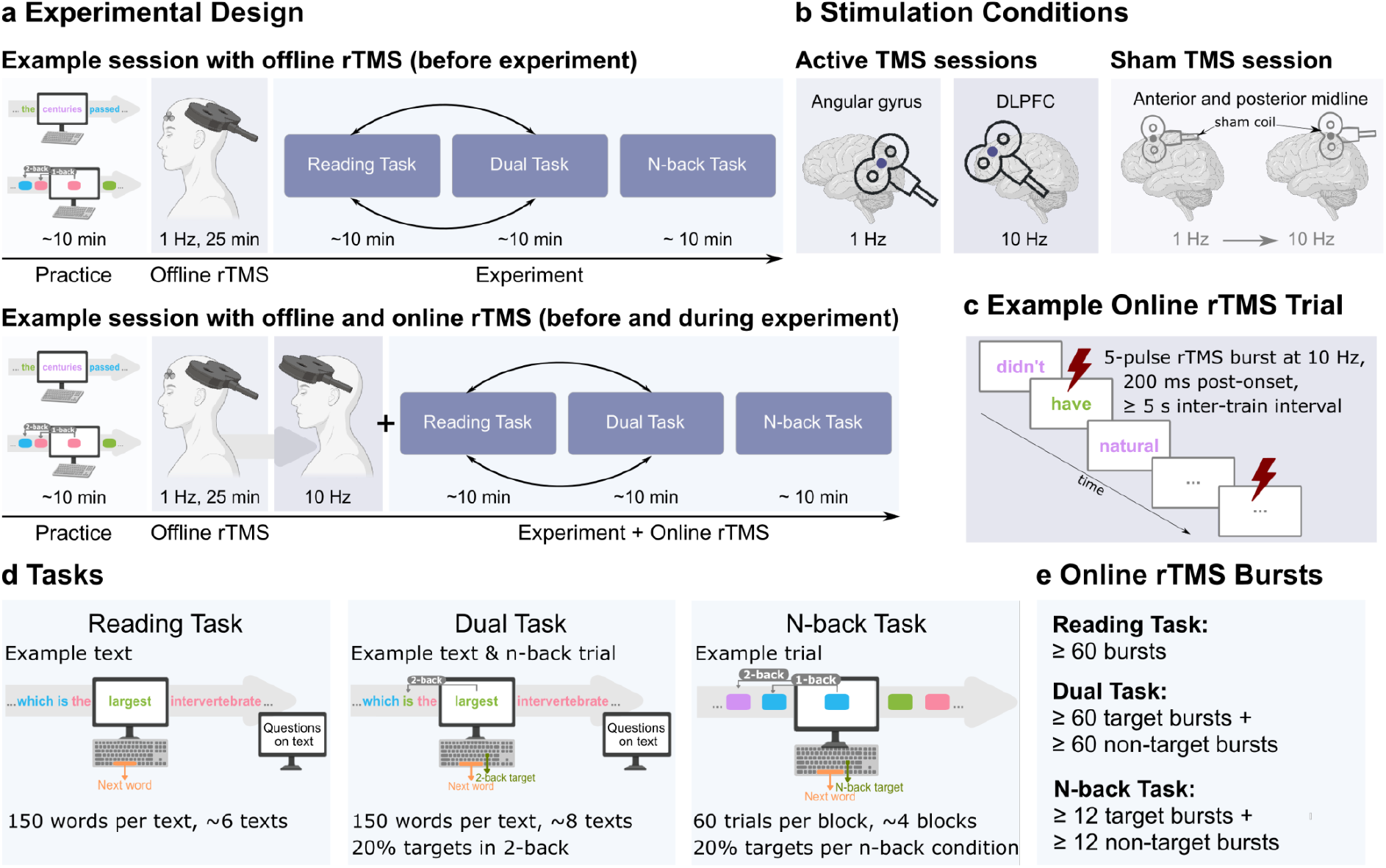
Experimental Design. **a**: Participants completed three sessions: an Offline rTMS, an Offline+Online rTMS, and a sham rTMS session. The sham session mimicked the Offline+Online session using a sham coil and different targets. The order of the reading and dual task was counterbalanced across participants. **b**: Stimulation targets for Active and Sham rTMS sessions. **c**: Example of an Online rTMS trial during reading. **d**: Example trials for the three experimental tasks. Texts were unique for each session and the presentation was balanced across sessions and participants. **e**: Minimum number of online rTMS bursts per task.

We hypothesized that stimulation effects on reading and working memory would be task-dependent, with larger effects under higher cognitive load (that is, either in a dual-task or a 2-back setting) than in single-task conditions. We predicted that AG stimulation would alter the effect of surprisal on reading times preferentially under concurrent working-memory demand, while DLPFC stimulation would modulate both the surprisal effect and working-memory performance, consistent with the proposed role of executive resources in processing prediction errors. By combining network-targeted stimulation, computational modeling, and task-load manipulation, this study provides a mechanistic account of when and how the neural systems supporting linguistic prediction and executive control interact during reading, thereby offering broader insight into how domain-general and domain-specific systems jointly support predictive processing under naturalistic conditions.

## Results

In this preregistered study (10.23668/psycharchives.16408), we report data from 25 healthy young participants who completed three experimental sessions: one session with 1 Hz rTMS over left AG before the experiment (Offline TMS), one session with 1 Hz rTMS over left AG before and 10 Hz rTMS over left DLPFC during the experiment (Offline+Online TMS), and one placebo sham TMS session (Sham TMS; Fig. 1a–c). In each session, participants performed three experimental tasks: a reading task, an n-back task (1-back and 2-back) assessing the color of squared rectangles, and a dual task combining reading with a concurrent 2-back paradigm (Fig. 1d). Using linear and non-linear mixed-effects regression, we examined effects of rTMS on response times and accuracy across tasks and explored interactions with task load (single n-back vs. dual task) and lexical surprisal as a proxy for predictive language processing during reading. We further tested whether rTMS influenced the underlying decision-making processes in n-back trials using hierarchical Bayesian drift-diffusion (DDM) models, which allowed us to jointly model reaction time and accuracy. Finally, we examined whether differences in the induced electrical field (E-field) strength predicted individual responses to TMS.

### TMS effects on reading increase in dual-task context

First, we asked if the stimulation conditions differentially modulated behavior with linear and nonlinear mixed effects models, which allowed us to control for a large number of known confounding variables, such as psycholinguistic estimates (e.g., word length and frequency), individual working memory abilities, and measured side effects during TMS. While the linear mixed model (LMM) for reading times revealed no main effect of TMS (F(2, 26) = 1.49, p = .243), a significant interaction between TMS and task (F(2, 102175) = 55.60, p < .001) and follow-up contrasts indicated that stimulation effects were task-dependent, with stronger speeding effects under dual-task load than during reading alone (Fig. 2a, b). Specifically, the speeding effect of Offline TMS relative to Sham was stronger in the dual task than during reading alone: Estimated reading times were 26.6 ms faster than under Sham in the dual task, compared with only 5.8 ms faster during reading alone (ratio = 0.981, *p* < .001). This task dependence was even more pronounced for Offline+Online TMS: Relative to Sham, estimated reading times were 18 ms faster in the dual task but 20.3 ms slower during reading alone (ratio = 0.946, *p* < .001). Thus, TMS effects were stronger in the dual task, shifting reading times toward faster responses relative to Sham under increased cognitive load. Moreover, within the reading task, a simple-effect analysis revealed that Offline+Online TMS slowed reading relative to Offline TMS (ΔRT = 26.08 ms, 95% CI [3.20, 48.95], p = 0.025), whereas this contrast was not significant in the dual task (Fig. 2b). The model further showed significant effects of session, task, trigger, n-back button press, and punctuation on reading times (Table S4).

**Figure 2.**
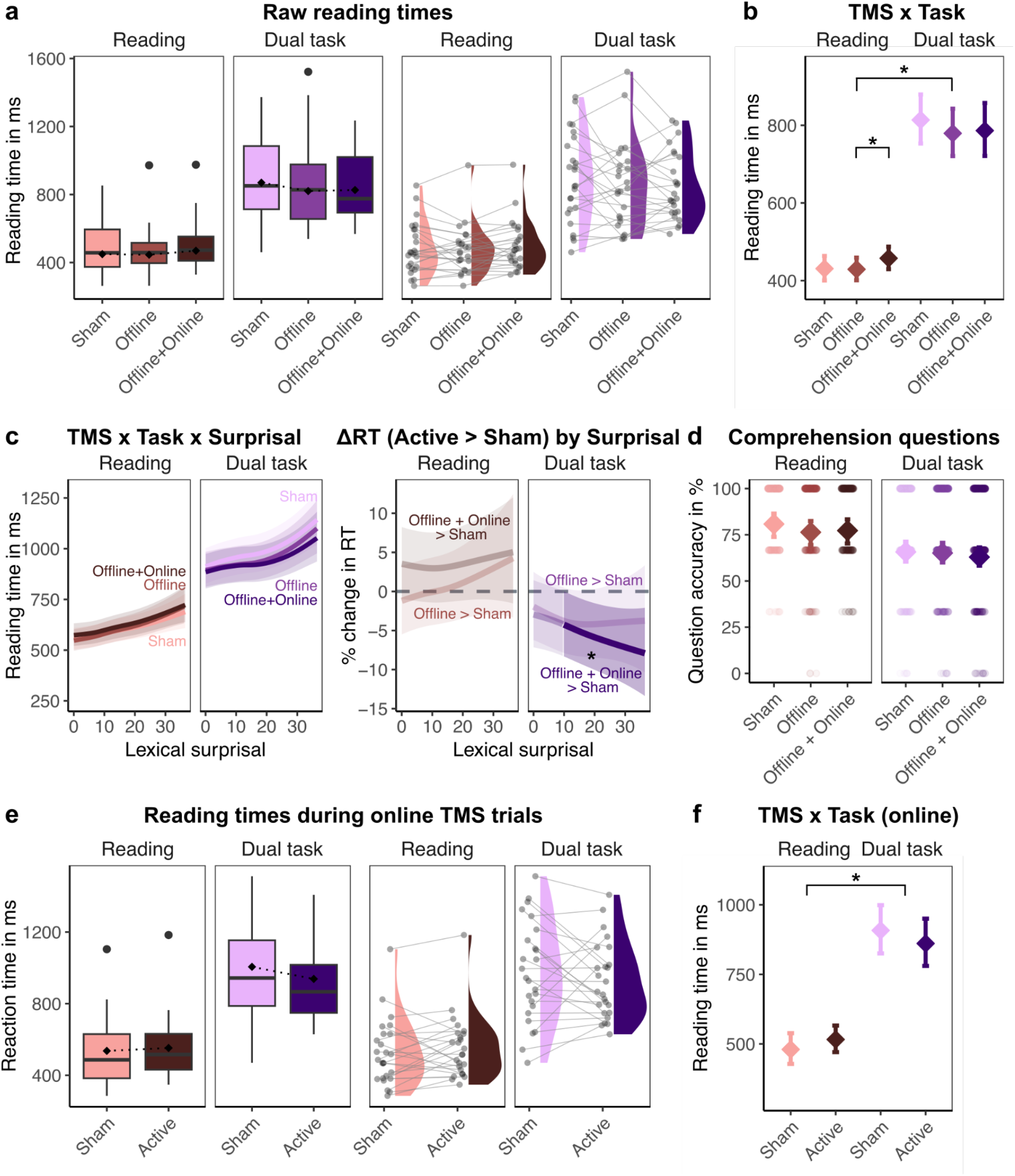
TMS effects on reading performance are stronger during dual than single task. During dual task, TMS reduces reading-costs of unexpected words. a: Raw reading times aggregated by stimulation condition (box plots) and distribution of raw data (half-violin plots) with means per participants as points. b: Estimated conditional means from the LMM for the TMS-by-task interaction. c: Estimated conditional smooths from the GAMM for the TMS-by-surprisal interaction on the left and percent changes in reading times (RT) relative to Sham TMS for each task and stimulation condition. Significant effects are opaque and marked with an asterisk. d: Raw aggregated data for accuracy in comprehension questions answered after each text. e: Raw reading times for online TMS trials only, aggregated by stimulation condition (box plots) and distribution of raw data (half-violin plots) with means per participants as points. f: Estimated conditional means from the LMM for the TMS-by-task interaction. Significant effects are opaque and marked with an asterisk.

A generalized additive mixed model (GAMM) examined potential nonlinear effects of lexical surprisal and time from task onset on reading times, including interactions between TMS, task, and lexical surprisal, as well as time progress. The model revealed a significant three-way interaction between surprisal, stimulation condition, and task (edf = 5.07, F = 5.66, p = .044). Follow-up contrasts indicated that only in the dual task, Offline+Online stimulation attenuated the positive association between surprisal and reading time relative to Sham, such that reading times increased less steeply as surprisal increased (Fig. 2c). No corresponding difference was evident in the reading task.

Over the duration of the experimental sessions, the GAMM showed an interaction between TMS and time (edf = 11.31, F = 3.15, p < .001). Follow-up contrasts showed that as time progressed, Offline TMS was associated with moderate speeding of reading times relative to Sham TMS (Table S4, Fig. S1a). The GAMM further revealed significant nonlinear effects of several psycholinguistic predictors (entropy, word length, frequency), the reading time of the previous word, and the surprisal of the previous word, confirming their influence on reading times (Table S4). Significant interactions also confirmed the task-dependent effects of surprisal and time progress (Table S4).

A separate analysis restricted to trials with 10 Hz online TMS (Active online vs. Sham online) again showed a significant interaction of TMS and task (F(1, 8923) = 29.06, p < .001): Similar to the previous analysis, Active online TMS speeded up reading in the dual task but not during the reading task alone (β_active − sham:dual − reading_ = -0.12, p < .001; Fig. 2e & f, Table S5). The GAMM further revealed a significant nonlinear three-way interaction between TMS, task, and time progress (edf = 2.00, F = 21.97, p < .001): Follow-up contrasts showed that during reading alone Active online TMS slowed reading more than Sham online, especially at the start of the task (Fig. S1b).

A generalized linear mixed model (GLMM) tested TMS effects on comprehension accuracy in both tasks (Fig. 2d). There was no main effect of TMS (*χ*^2^(2) = 1.63, p = .442) and no TMS-by-task interaction (*χ*^2^(2) = 0.68, p = .712). Task, however, had a significant effect (*χ*^2^(1) = 36.52, p < .001), with poorer accuracy during the dual task (β_dual task - reading_ = −0.79, p < .001). Moreover, older age was associated with poorer accuracy (*χ*^2^(1) = 6.99, p = .008; β = −0.04, p = .029), whereas better executive abilities were associated with higher accuracy (*χ*^2^(1) = 4.74, p = .029; β = 0.24, p = .029; see Table S6 for full model results).

Together, the linear and nonlinear analyses showed that cognitive load shaped the magnitude and direction of TMS effects: Stimulation generally facilitated reading under dual-task demands but had weaker effects during reading alone. Importantly, these task-dependent changes in reading speed occurred without compromising comprehension accuracy.

### TMS preferentially alters working-memory performance under high load

After establishing robust TMS effects on reading performance, we zoomed in stimulation effects on working memory across different cognitive loads. Specifically, we investigated TMS effects on correct n-back reaction times during the single n-back (1- and 2-back) and the 2-back dual task (Fig. 3a). While the LMM showed no significant main effect of TMS (F(2, 584) = 2.14, p = .119), there was a significant interaction between TMS and task (F(4, 10305) = 2.42, p = .046; Fig. 3b). Follow-up contrasts showed that for all tasks, Offline+Online TMS was linked to faster reactions than Sham, although only significant in the 2-back condition (ΔRT = 78.05 ms, 95% CI [5.31, 150.79]). The model further showed significant effects of session, task, trigger, executive abilities, years of education, and side effects on reading times (Table S7).

**Figure 3.**
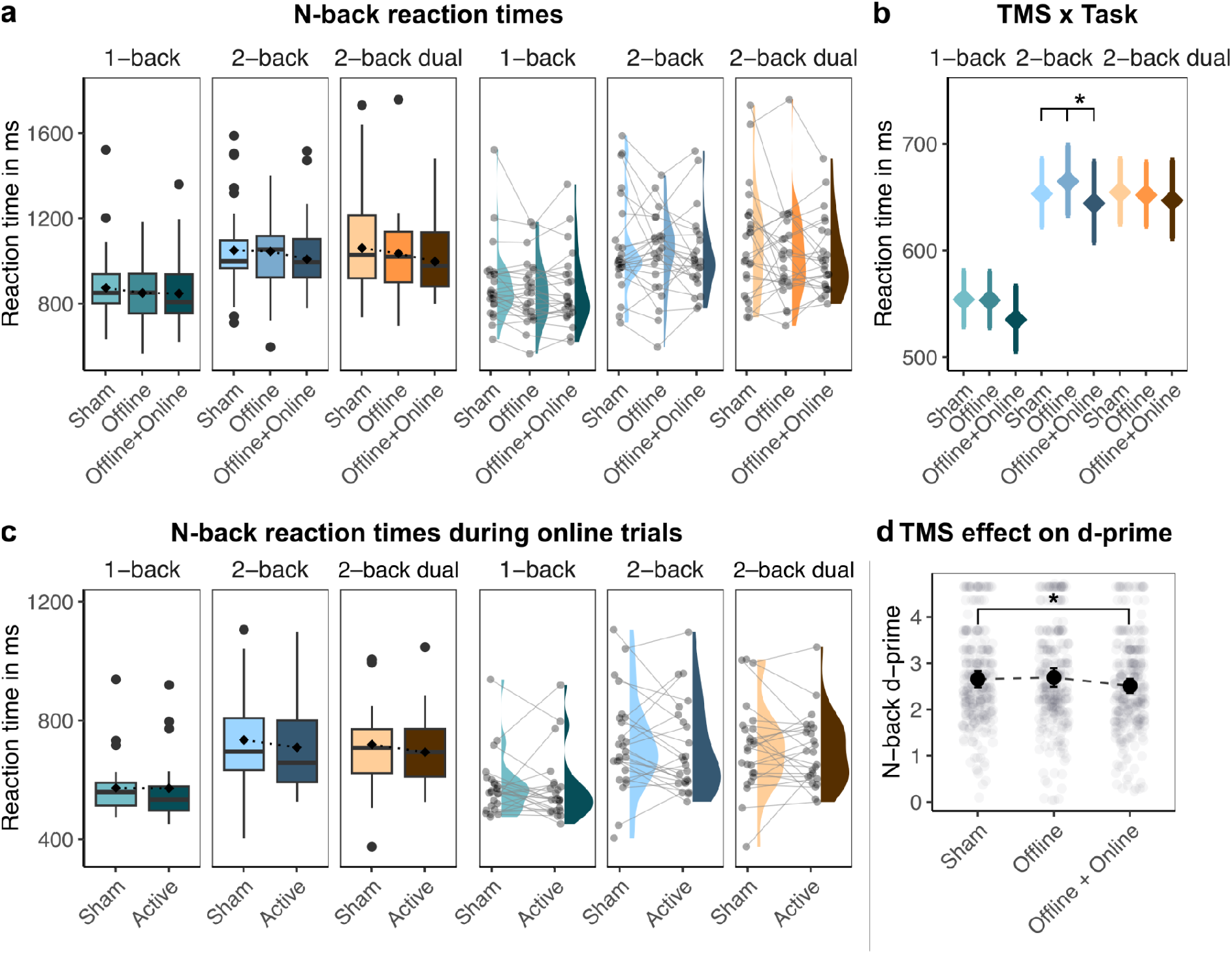
TMS effects on working memory performance are strongest during more demanding 2-back condition. **a**: Raw reaction times aggregated by stimulation condition (box plots) and distribution (half-violin plots) with means per participants as points. **b**: Estimated conditional means from the LMM for the TMS-by-task interaction. **c**: Raw reaction times for online TMS trials only, aggregated by stimulation condition (box plots) and distribution of raw data (half-violin plots) with means per participants as points. **d**: Estimated conditional means from the LMM for the TMS main effect on d-prime values. Significant changes at p < .05 relative to Sham are marked with an asterisk.

A GAMM tested nonlinear interactions of TMS, task, and time progress. There was no significant interaction between TMS, task, and time (edf = 8.62, F = 1.44, p = .164) but a significant interaction between task and time progress (edf = 4.92, F = 22.64, p < .001, Table S7), revealing slightly slower reaction times for the 2-back and 2-back dual task compared with the 1-back task over time.

Separate models were fit to trials with 10 Hz online TMS (active online vs. sham, Fig. 3c). Results from the LMM showed no significant main effect of TMS (F(1, 1517) = 0.57, p = .451) or interactions of TMS and task (F(2, 2599) = 1.46, p = .233) but significant effects of task, session, executive abilities, and years of education (Table S8). Similarly, the GAMM did not show an interaction of TMS conditions with time progress restricted to the online TMS trials (Table S8).

An LMM testing TMS effects on d-prime in all tasks showed a significant main effect of TMS (F(2, 45) = 6.36, p = .004; Fig. 3d). The model output showed that d-prime significantly decreased after Offline+Online TMS (β = -0.14, p = .011) relative to Sham TMS. Moreover, results revealed significant effects of task, executive abilities, the block number of the experiment, and reading span (see Table S6).

In summary, matching the load-dependent effects on reading, TMS effects on working memory were most evident under high cognitive demand: Offline+Online TMS accelerated correct responses, most clearly in the 2-back task, but reduced overall response sensitivity.

### Computational modeling of n-back decisions in n-back and dual-task contexts

To better understand the effects of stimulation and surprisal on the decision-making process underlying n-back responses, we fitted hierarchical Bayesian DDMs jointly modeling reaction times and accuracy. Here, we focused on the n-back trials (single n-back and dual tasks), as reading-only did not yield trial-wise accuracy data. Separate models were fitted to n-back responses in the single n-back conditions and in the dual task, allowing the inclusion of additional psycholinguistic predictors, such as lexical surprisal in the latter. Model comparison evaluated alternative specifications of parameter modulation, including TMS- and task-related effects on the four core DDM parameters: drift rate *v*, boundary separation *a*, non-decision time *t*, and starting point bias *z* (Fig. 4a, see Table S3 for full model formulas). For both datasets, comparison based on leave-one-out cross-validation favored models in which drift rate *v* and decision boundary *a* included TMS-by-task interactions as fixed effects with by-participant random slopes for TMS and task (Fig. 4b,c), suggesting that these parameters best capture key aspects of the participants’ choices and response times. We report mean posterior estimates with 95% highest density intervals (HDIs) and their probability of direction (pd). The pd ranges from 50% to 100% and reflects the probability that a parameter is strictly positive or negative, thereby indexing the existence of an effect [30,31].

**Figure 4.**
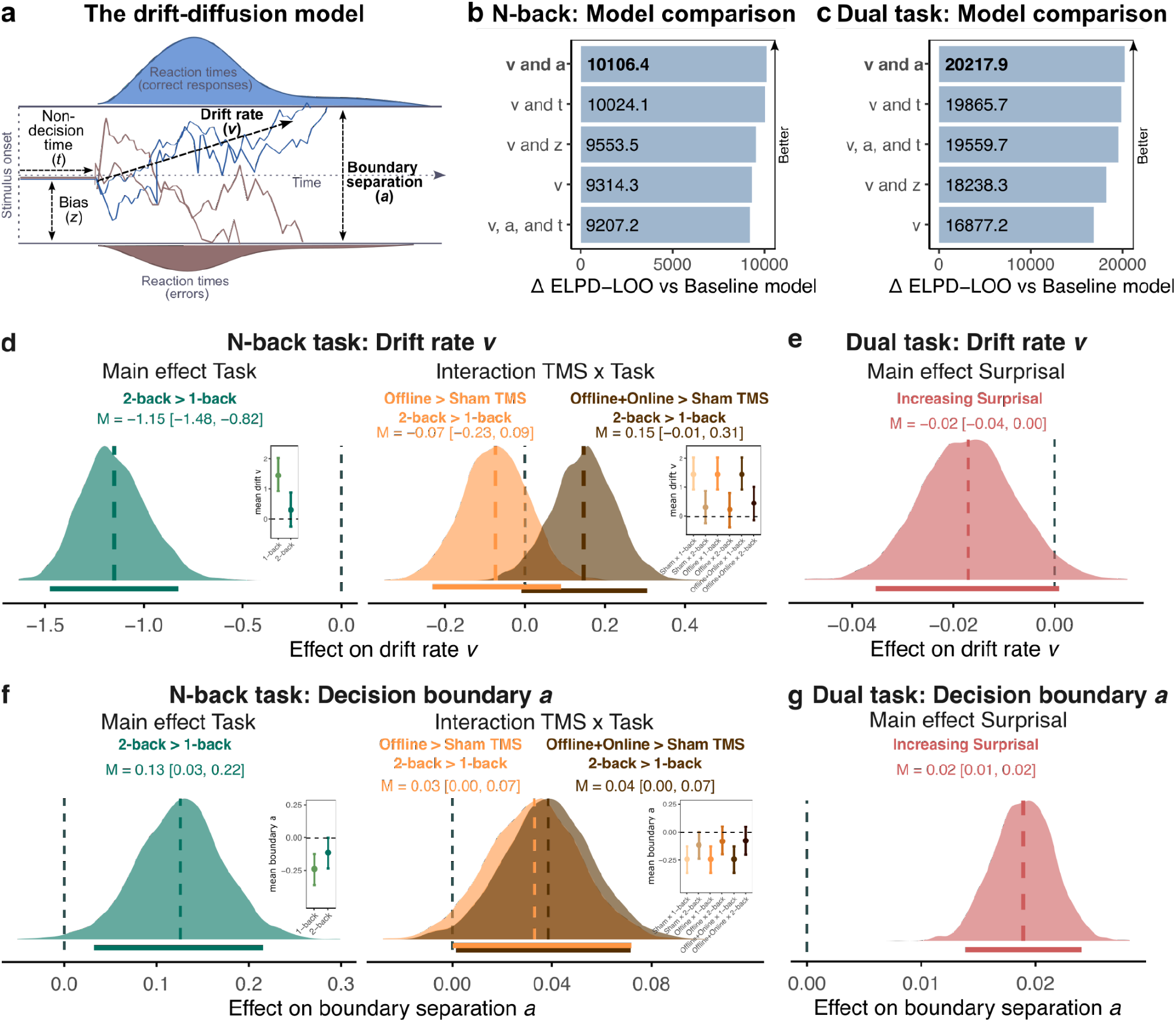
Computational modeling of TMS effects on n-back performance: Drift rate and boundary separation are affected by TMS. **a**: Illustration of the drift-diffusion model and its core parameters. Favored parameters by model comparison are marked in bold. **b** and **c**: Results of model comparisons for n-back and dual task, respectively, via leave-one-out (LOO) cross-validation. Values are differences in expected log predictive density (ELPD) against a baseline model with no parameter modulated. Larger values indicate better estimated out-of-sample predictive performance. **d**: Predicted posterior estimates for effects in n-back task modulating drift rate. Since distributions show differences between factor levels, the insets display the recovered estimates for individual factor levels. **e**: Predicted posterior estimate for the effect of lexical surprisal on drift rate in dual task trials. **f**: Predicted posterior estimates for effects in the n-back task modulating boundary separation. Since distributions show differences between factor levels, the insets display the recovered estimates for individual factor levels. **g**: Predicted posterior estimate for the effect of lexical surprisal on boundary separation in dual task trials.

### TMS facilitates n-back performance under high task load

For n-back reactions, posterior estimates indicated no main effect of TMS on drift rate (v_Offline - Sham_ = −0.04, [−0.21, 0.13], pd = 69%; v_Offline+Online - Sham_ = −0.06, [−0.24, 0.13], pd = 75%; Supplementary Figure S2a). There was a strong task effect, with lower drift rates in the 2-back relative to the 1-back task (v_2-back - 1-back_ = −1.15, [−1.48, -0.82], pd = 100%; Fig. 4d). The TMS-by-task interaction suggested a stronger effect of Offline+Online TMS in the 2-back than in the 1-back task (v_Offline+Online - Sham:2-back - 1-back_ = 0.15, [−0.009, 0.31], pd = 96%). Posterior estimates showed a higher drift rate for the 2-back compared with the 1-back task during Offline+Online TMS (Fig. 4d). This effect was not present for Offline TMS (v_Offline - Sham:2-back - 1-back_ = -0.07, [−0.23, 0.09], pd = 81%). Additional effects indicated reduced drift rates during trials with online TMS and increased drift rates in sessions 2 and 3 relative to session 1. There was no evidence for a systematic effect of time on drift rate (see Table S9 and Fig. S2a for full estimates).

Similar results emerged for boundary separation *a*. There was no main effect of TMS (a_Offline - Sham_ = 0.00, [−0.08, 0.07], pd = 50%; a_Offline+Online - Sham_ = −0.05, [−0.14, 0.03], pd = 89%;

Supplementary Figure S2b), but a strong task effect with increased boundary separation in the 2-back than the 1-back task (a_2-back - 1-back_ = 0.13, [0.03, 0.22], pd = 99%; Fig. 4f). The TMS-by-task interaction indicated moderately larger boundary separation in the 2-back relative to the 1-back task under both stimulation conditions (a_Offline - Sham:2-back - 1-back_ = 0.03, [0.00, 0.07], pd = 97%; a_Offline+Online - Sham:2-back - 1-back_ = 0.04, [0.002, 0.07], pd = 98%; Fig. 4f), indicating increased boundaries after either effective stimulation. Additional effects showed increasing boundary separation over time, as well as reduced boundary separation in later sessions and during trials with online TMS (Table S9 and Fig. S2b for full estimates).

### Lexical surprisal propagates into concurrent working-memory decisions

In the dual task, posterior estimates indicated no main effect of TMS on either drift rate (v_offlineOffline - shamSham_ = −0.05, [−0.15, 0.05], pd = 83%; v_offlineOffline+onlineOnline - shamSham_ = 0.03, [−0.05, 0.11], pd = 79%; Fig. S3a) or boundary separation (a_offlineOffline - shamSham_ = −0.05, [−0.13, 0.04], pd = 89%; a_offlineOffline+onlineOnline - shamSham_ = 0.008, [−0.07, 0.08], pd = 59%; Fig. S3b). There was a moderate effect of lexical surprisal on drift rate, which decreased with increasing surprisal (v = −0.02, [−0.04, 0.0008], pd = 97%; Fig. 4e). In contrast, surprisal showed a stronger effect on boundary separation, which increased with higher surprisal (a = 0.02, [0.01, 0.02], pd = 100%; Fig. 4g). The effects of surprisal on drift rate and boundary separation were not credibly modulated by TMS (see Table S10).

The model also included additional psycholinguistic predictors including word length, frequency, and entropy, which showed stronger associations with boundary separation than with drift rate. Moreover, drift rate decreased over time, and during online TMS trials, but increased in later sessions. In contrast, boundary separation decreased over time, in later sessions, and during online TMS trials (see Table S10 and Fig. S3 for full estimates).

In summary, the DDMs paralleled the behavioral finding that TMS effects were most pronounced under high cognitive load, revealing load-dependent changes in evidence accumulation and response caution. They additionally uncovered cross-task spillover: More surprising words slowed evidence accumulation and increased response caution in concurrent n-back decisions.

### TMS-induced effects scale with individual E-field strength

Given potential dose-response relationships of the cortical stimulation strength, we performed post-hoc simulations to characterize the TMS-induced electric field strength at our target regions and included individual E-field values as predictors in regression models of behavioral performance. To this end, we calculated task-specific LMMs for each target region (AG and DLPFC). E-fields showed local stimulation maxima at the intended coordinates in left AG and DLPFC with stimulation extending into the posterior superior and middle temporal gyrus in case of the AG and the inferior frontal gyrus (pars triangularis) for the DLPFC (Fig. 5a).

**Figure 5.**
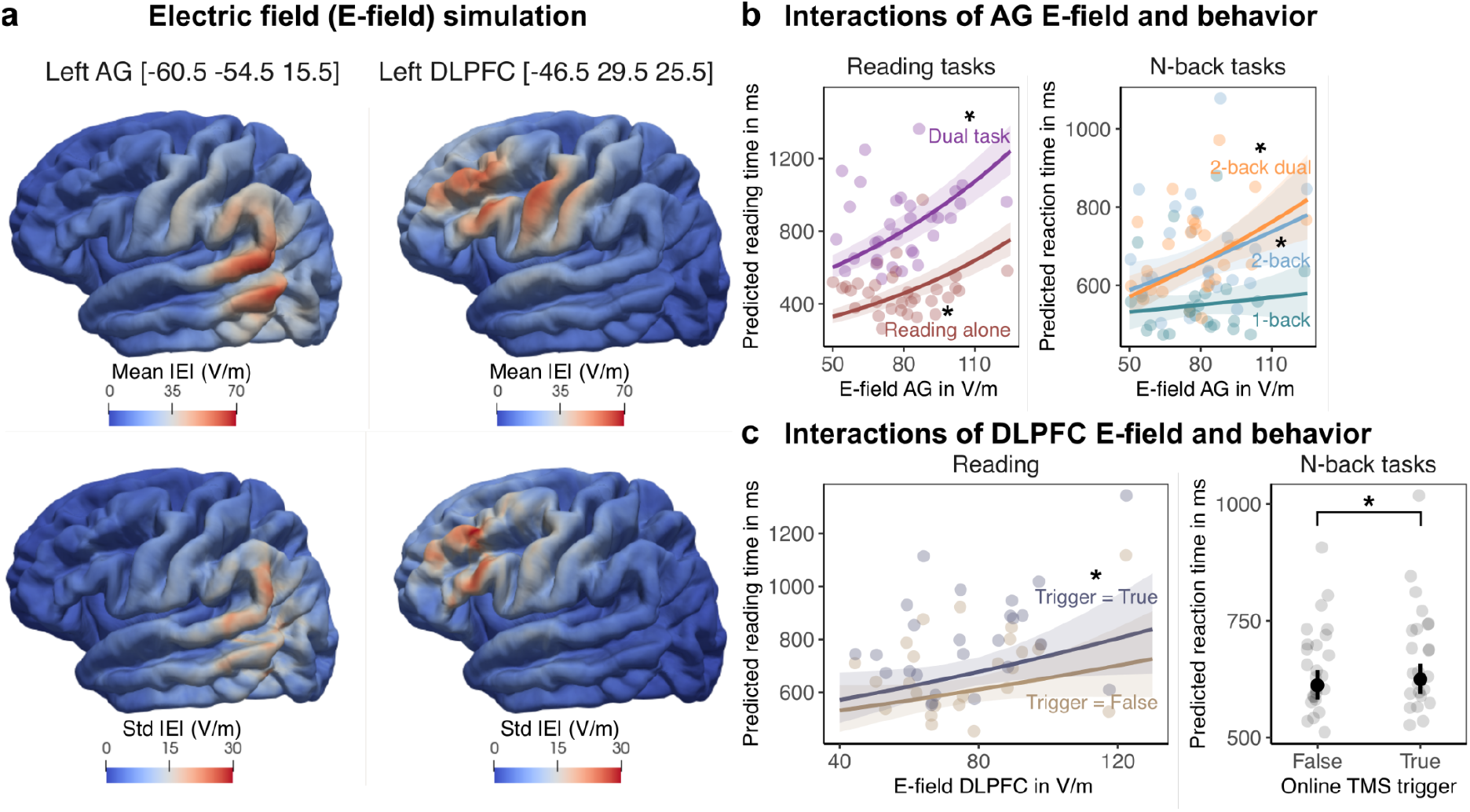
E-field strength predicts TMS effects on behavior. **a**: Post-hoc simulated electric fields (E-fields) at target MNI coordinates. **b**: Significant interactions between individual E-field strength in the AG and reading and n-back reaction times. **c**: Significant interaction between E-field strength in the DLPFC and reading times. Points show individual effects and significant effects at p < .05 are marked with an asterisk.

The E-field strength in the AG had a main effect on reading times (F(1, 60903) = 1282.31, p < .001; Table S11) and showed an interaction with task (F(1, 11370) = 5.75, p = .016). Results showed that larger E-field strength in the AG was associated with slower reading and this effect was slightly stronger in reading alone relative to the dual task (β_reading – dual task_ = -0.02, p = .016; Fig. 5b).

Similarly, for the n-back tasks, larger AG E-field strength was linked to overall slower reaction times (F(1, 315) = 22.84, p < .001; Table S12) and this effect was additionally modulated by task (F(2, 36) = 6.97, p = .003). Specifically, both the single 2-back and 2-back dual tasks were more strongly affected by greater E-field strength relative to the single 1-back task.

Since the DLPFC was only stimulated in the Offline+Online session during selected trials, we investigated potential effects of E-field strength specific to this session. For reading times, the E-field in the DLPFC showed an interaction with TMS train, thus whether there was an actual online stimulation on the trial or not (F(1, 36634) = 14.53, p < .001; Table S11). With increasing E-field strength, the effect of an online TMS train increased, leading to slower reading time in that trial (β = 0.0008, p < .001; Fig. 5c). For n-back reaction times, no similar effect emerged (Table S12).

## Discussion

The present study tested whether the neural systems supporting linguistic prediction during reading interact with executive resources as cognitive demands increase. By combining naturalistic reading with a concurrent nonverbal working-memory task, network-targeted rTMS, and computational modeling, we identified three converging findings. First, stimulation effects on reading were strongly task-dependent, accelerating reading times under dual-task load but not during reading alone. Combined stimulation of left AG and DLPFC also attenuated the effect of lexical surprisal selectively during dual-task reading, whereas it did not modulate the surprisal effect during reading alone. Second, TMS altered working-memory performance and its underlying decision dynamics most clearly under high task load. Third, during the dual task, surprisal affected performance beyond reading itself: Unexpected words reduced the efficiency of evidence accumulation and increased response caution in the concurrent n-back task. Together, these findings provide evidence that contextual prediction during reading is dynamically coordinated with domain-general executive control. Rather than operating as an isolated property of the language system, the behavioral impact of prediction depends on how cognitive resources are allocated across concurrent goals.

### Load-specific modulation of lexical surprisal

The load-specific modulation of lexical surprisal provides direct evidence for an interaction between language and executive systems. Reading times normally increase with surprisal because less expected words require greater updating of the current linguistic representation [4–6]. In the present study, combined AG and DLPFC stimulation flattened this relationship selectively during the dual task: Highly surprising words produced a smaller increase in reading time than under sham TMS. AG stimulation alone produced a similar but weaker attenuation, suggesting that perturbing AG-supported contextual representations may already reduce the influence of lexical predictions, while additional DLPFC stimulation amplifies this effect when executive resources are taxed. This was not a general change in reading speed or lexical sensitivity, because the same modulation did not emerge during reading alone. Taken together, these results indicate that stimulation changed how strongly contextual expectations shaped moment-to-moment behavior when readers also had to maintain and update a nonverbal working-memory sequence. Importantly, these changes occurred without an additional stimulation-related reduction in comprehension accuracy, arguing against a general disruption of text processing.

This finding aligns with a growing body of behavioral, eye-tracking, and electrophysiological evidence showing that surprisal effects are attenuated under concurrent cognitive demands, including simultaneous memory tasks [9,10,32], increased visuospatial memory load [8], diverted attention [33,34], and rapid presentation of speech [35,36]. Here, we extend these findings by identifying a neural system through which this attenuation can be modulated. The surprisal effect was attenuated only during dual-task reading and only following combined AG and DLPFC stimulation, not AG stimulation alone. This pattern reinforces the notion that unexpected words incur processing costs [37], while showing that the magnitude of this cost depends on the executive resources available to maintain and update context-derived expectations when resources are shared across concurrent goals.

The finding also stands in contrast with accounts that separate linguistic prediction from domain-general executive control [11,38,39]. Although prediction may proceed largely within the language system during reading alone, maintaining precise contextual expectations appears to recruit executive control when another task competes for cognitive resources. Our results therefore identify cognitive competition as a boundary condition under which linguistic prediction becomes dependent on domain-general control.

### Bidirectional interaction of predictive language and executive systems

Complementary evidence for the interaction of language and executive systems came from the dual-task drift-diffusion model. Although n-back judgments concerned the words’ font color rather than their linguistic content, lexical surprisal influenced the dynamics of the concurrent working-memory decision. As surprisal increased, evidence accumulation became less efficient and participants adopted more cautious response behavior, reflected in lower drift rates and greater boundary separation. Thus, the processing cost of an unexpected word was not confined to reading but extended to a simultaneous working-memory decision of the same stimulus [8,9]. This provides a computational account of dual-task interference: Updating the linguistic representation competes with accumulating relevant evidence for the n-back decision process, while the increased decision boundary may reflect compensation for this interference. Together with the reading-time results, these findings reveal a bidirectional interaction. Working-memory demands regulate the behavioral impact of linguistic prediction, while prediction errors alter the dynamics of concurrent working-memory decisions.

TMS altered working-memory performance and its underlying decision dynamics most clearly under high task load.

#### Load-specific modulation of non-verbal working memory

The stimulation results on working-memory performance during the n-back task further support a load-dependent effect of TMS. Combined AG and DLPFC stimulation produced faster responses most clearly in the demanding 2-back condition, and the drift-diffusion model showed that it attenuated the load-related reduction in drift rate while raising boundary separation. The concurrent increase in drift rate and boundary separation indicates an adaptive reconfiguration of decision-making. Stimulation improved the efficiency of task-relevant evidence accumulation while increasing the amount of evidence required for a decision, potentially in response to greater uncertainty or interference in the demanding 2-back condition. This beneficial TMS effect on working memory is in line with previous work and meta-analyses on high-frequency rTMS of DLPFC [28,29]. Together, these results show that combined TMS of AG and DLPFC shifted the balance between distinct components of working-memory decisions, with the clearest effects emerging when monitoring demands were high. This parallels the reading results: In both domains, stimulation effects became most relevant when the system had to handle greater cognitive load.

The reduction in d-prime after combined AG and DLPFC TMS provides a complementary block-level perspective on this shift. Although d-prime summarizes overall discrimination, it collapses across trial-level information. The DDM instead jointly models individual choices and response times, separating the efficiency of evidence accumulation from the amount of evidence required for a response. A lower aggregate d-prime therefore does not contradict a relative increase in drift rate on demanding trials. Rather, the combined results show that stimulation did not uniformly improve or impair working memory, but changed the mechanisms through which decisions were reached. This distinction also illustrates the value of process-level modeling of TMS effects on cognition, providing insights into stimulation-induced mechanistic changes [40–42].

### Limitations and outlook

The weaker modulation of surprisal after AG stimulation alone was unexpected given the region’s established role in semantic integration [14–18]. Although the effect was descriptively in the same direction as after combined stimulation, it did not reliably differ from sham TMS. Importantly, however, the E-field analyses indicate that AG stimulation was behaviorally effective: Stronger stimulation strength at the target region was associated with slower reading and n-back responses, with larger effects in the more demanding n-back conditions. Thus, AG perturbation produced dose-sensitive processing costs, but its influence on surprisal became pronounced only when DLPFC stimulation was added under competing task demands. This pattern suggests that constructing and maintaining a contextual representation is not identical to determining how strongly prediction error influences moment-to-moment behavior [16,43,44]. The AG may support the integration of incoming words with the preceding context, whereas DLPFC-mediated control becomes important for maintaining and prioritizing that context when another task competes for resources. The stronger surprisal effect after combined stimulation therefore points to load-dependent prediction as a network-level function rather than a property of AG alone.

Our findings open several important avenues for future work. Because online DLPFC stimulation was administered after offline AG stimulation, the combined condition identifies a contribution of both regions but cannot isolate the independent role of DLPFC. A design involving a separate DLPFC-only condition will be required to determine whether the contributions of AG and DLPFC are additive or interactive. Further, more temporally resolved approaches such as chronometric TMS could determine whether the attenuated surprisal effect reflects reduced weighting of contextual predictions or faster resolution of prediction errors, providing further insight into the mechanistic system of predictive language processing.

## Conclusion

In conclusion, by combining naturalistic reading, neurostimulation, and computational modeling, this study reveals how language and executive systems interact under the competing demands characteristic of everyday cognition. Our findings identify cognitive competition as a key condition under which linguistic prediction becomes coupled to executive control. Combined AG and DLPFC stimulation altered the effect of lexical surprisal only when reading competed with working-memory monitoring, while lexical surprisal affected a concurrent n-back decision process. This pattern is consistent with complementary roles of AG and DLPFC: The AG supports contextual representations from which linguistic expectations arise, whereas the DLPFC helps maintain and prioritize those representations when multiple goals compete. Together, these findings show that linguistic prediction is embedded in a broader cognitive system that dynamically coordinates language and executive control under competing demands.

## Materials and Methods

### Participants

Twenty-five healthy younger adults (M = 31.3, SD = 6.6, range = 21–41 years; 13 female) participated in the study. The sample size was determined based on an a priori power analysis with G*Power 3.1 [45] for a within-subject, repeated-measures design, indicating that at least 23 participants were required to detect a medium effect size (Cohen’s d = 0.564) with a Type I error probability of 0.05 and a power of 90%. To account for potential data loss due to technical issues during acquisition, we included 25 participants.

All participants were right-handed, native German speakers with no contraindications to MRI or TMS, and no colour blindness as assessed by the Ishihara Test [46]. After each session, participants completed a questionnaire on TMS side effects [47]. Additionally, at the end of the sham TMS session, neuropsychological profiles were assessed using the Trail Making Test [48], Digit Symbol Substitution Test [49], and the Reading Span Test [50]. Participants were informed about the experimental procedures but were blinded to the different TMS conditions and their order. The study was approved by the local ethics committee of the Medical Faculty at Leipzig University and conducted in accordance with the Declaration of Helsinki. Written informed consent was obtained from all participants before the experiment.

### Design

Participants completed three experimental sessions of a within-subject repeated-measures design, which were separated by at least 5 days. At the beginning of each session, participants (re-)familiarized themselves with the tasks. They then received offline low-frequency rTMS over left AG and subsequently performed the experiment (Fig. 1a and b). During the offline+online session, they additionally received high-frequency rTMS over the left DLPFC during the experiment. The sham stimulation session mimicked the offline+online session using a sham TMS coil. The order of the stimulation conditions was counterbalanced across participants.

### Tasks and stimuli

Participants performed three experimental tasks: a reading task, an n-back task (1-back and 2-back) assessing the colour of squared rectangles, and a dual task that combined reading with the 2-back paradigm (Fig. 1d). Task paradigms were adapted from a previous, similar study without TMS in our laboratory [10].

For both reading and dual tasks, participants read 150-word newspaper articles, pressing the space bar with their left hand to advance to the next word, minimizing TMS-induced interference with motor components during reading and online stimulation. Articles were adapted from a freely available newspaper archive ("Der Spiegel"), standardized for readability (Wiener Sachtextformel 7 – 8.5), length, and screened for neutral content. A total of 48 articles were prepared, of which 16 were randomly assigned per session and participant, without repetition. After each text, three multiple-choice questions assessed comprehension. In the dual task, participants performed the reading task while simultaneously completing a 2-back task on the font color of each word, pressing a button with their right hand whenever the current word’s color matched that of the word two positions earlier.

For the n-back task, participants performed 1-back and 2-back tasks with colored rectangles. In the 1-back condition, they pressed a button when the current rectangle’s color matched the preceding one; in the 2-back condition, they responded when the color matched that of the rectangle two positions prior. All tasks were self-paced. Session order and task assignment were counterbalanced across participants. Texts and stimuli were unique to each session and balanced for presentation order. The experiment was programmed and run in Psychopy v.2024.2.5 [51] in Python.

### Transcranial Magnetic Stimulation

In the offline TMS session, participants received 1 Hz offline rTMS at 100% resting motor threshold (rMT) for 25 minutes over the left AG before the experiment (see Fig. 1 a and b). During the offline+online session, participants again received 1 Hz rTMS over the left AG before the experiment, immediately followed by 10 Hz rTMS at 100% rMT over the left DLPFC during the experiment. To maximize effective blinding, the sham session mimicked the offline+online session using a sham TMS coil over two midline targets (anterior and posterior midline). At the end of each session, participants completed a computerized version of an established TMS side effects questionnaire [47]. The outcome was used in subsequent statistical analyses.

During offline rTMS, 1500 pulses were delivered in total. During online rTMS, 5-pulse trains were delivered 200 ms after trial onset, with a minimum inter-train interval of 5 s (Fig. 1c). A predefined minimum number of TMS trains per task ensured at least 228 trains (1140 pulses) per session (Fig. 1e). If the minimum number of trains for a given task was reached while participants were still completing a task block, additional trains (≥ 5 s apart) were delivered until the block ended. The distribution of TMS trains across tasks and stimulation conditions is shown in Figure S4a. In the dual and n-back tasks, we further ensured that TMS trains were equally distributed across target and non-target trials (Fig. S4b). To equalize session duration across all stimulation conditions, PsychoPy was programmed to count potential TMS bursts silently during the offline-only rTMS session. Average, minimum, and maximum session durations by condition are shown in Figure S5. No significant differences were found in the number of delivered TMS trains or overall session length between TMS conditions.

The rMT was assessed at the beginning of the first session and defined as the lowest stimulation intensity inducing motor evoked potentials of ≥ 50 µV at least 5/10 times in the relaxed first dorsal interosseous muscle when single-pulse TMS was applied to the left motor cortex. TMS was delivered with a figure-of-eight coil (MagVenture MCF-B65 for effective stimulation and MCF-P-B65 placebo coil for sham stimulation) connected to a MagPro X100 stimulator (MagVenture) and guided by stereotactic neuronavigation (TMS Navigator, Localite). To this end, the participant’s head was co-registered onto an individual structural MR scan taken from the in-house database or newly acquired at a 3T Siemens MR scanner.

For both AG and DLPFC stimulation, the coil was oriented at 45° (Fig. 1b). When this position was not mechanically feasible for AG stimulation because the coil holder was obstructed, the coil was rotated by 180° and the direction of current flow was reversed, thereby preserving the intended direction of the induced current relative to the cortical surface. For the midline targets during sham stimulation, the placebo coil was oriented at 0° with the handle pointing posteriorly. Target locations (in MNI space) for the left AG (x: -60.5, y: -54.5, z: 15.5) and DLPFC (x: -46.5, y: 29.5, z: 25.5) were derived from an unpublished fMRI study using the same experimental paradigm in a separate group of young healthy adults (n = 34, M = 29.8 ± 5.7 years, range: 19-39 years). See Supplementary Methods for details on MRI data acquisition, preprocessing and statistical analysis of fMRI data. The AG target was based on the contrast Reading only > Rest, whereas the DLPFC target was based on the contrast 2-back > 1-back to base the target on the contrast with the highest executive demand (Fig. S6, Tables S13 and S14). Target coordinates for the placebo stimulation using a sham TMS coil were chosen to represent one anterior and one posterior midline structure (x: 0, y: -35 z: 95; x: 0, y: 10, z: 90). The MNI coordinates were converted into participants’ native space and used for neuronavigation.

We performed post-hoc E-field simulations using SimNIBS v.4.6 [52] to characterize location, extent, and strength of the e-field induced by rTMS over the AG and DLPFC. E-field values were extracted by defining a region of interest with a 10 mm radius around the stimulation coordinates. From this region, values at the 95% percentile were extracted from the individually calculated E-fields, focusing exclusively on gray matter. To assess the effect of the induced stimulation strength over the target areas, we calculated task-specific LMMs for each ROI including the e-field values as predictors and behavioral performance as outcome measure (see Table S1 for model formulas).

### Data Analysis

Statistical models analysed per-word reading times (reading and dual-task reading) and per-trial reaction times in the n-back task (1- and 2-back task and dual task). With respect to performance accuracy, we analysed percent accuracy in the comprehension questions for the reading and dual tasks, and n-back accuracy in the single and dual n-back tasks. We fitted non-linear and linear mixed-effects regression models as described below.

### Preprocessing

Since participants performed self-paced versions of all tasks, reading and reaction times were measured via per-word and per-trial button presses. Excessive outlier values were excluded following a recommended procedure [53]. In short, for each participant, session, task, and task block, we calculated the Proportion of Maximum Scaling [POMS; 54] square-root transformed these values, z-transformed the result, and excluded observations with z-scores outside the range of −2 to 2 [10,53]. This procedure removed 1.5% (2,148) of all trials (142,750). To facilitate interpretability of units in the statistical analyses, we subsequently worked with the cleaned, log-transformed reading and reaction times.

For the analysis of comprehension question accuracy in the reading and the dual task blocks, we calculated the percent correct responses in the three comprehension questions per text.

For the analysis of accuracy in the n-back task, we calculated the sensitivity index detection-prime (d-prime) in the dual and single n-back tasks. We excluded task blocks with a d-prime ≤ 0 since a d-prime of 0 indicates performance at chance level, while a d-prime below 0 indicates that participants did not reliably perform the n-back task [55]. This criterion removed 3.6% (3,388) of all trials (94,109) of the dual and single n-back tasks.

### Generation of Word Surprisal and Entropy Scores

For all texts presented in the reading task, we computed word-level surprisal and entropy. Both measures capture lexical predictability, but at different stages. Word entropy reflects uncertainty about the upcoming word before it is encountered, whereas word surprisal reflects how unexpected the actually observed word is given the preceding context. Higher surprisal values correspond to lower probability and are often taken to index greater processing demand. Surprisal for word *w_n_* was defined as:

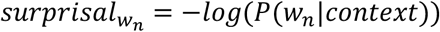

Word entropy was defined as the Shannon entropy of the predictive distribution over possible next words, which represents the average surprisal of all possible lexical continuations:

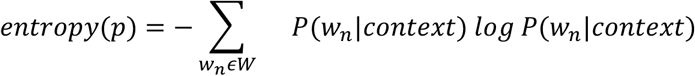

Both measures were derived from the probability distribution generated by a 12-layered GPT-2 model pretrained on German texts by the MDZ Digital Library team and the corresponding tokeniser, which are both available via the Huggingface model hub.

We included both measures because they capture complementary components of prediction. Entropy quantifies uncertainty in the distribution of possible continuations before a word appears, whereas surprisal quantifies the information conveyed by the continuation that is actually encountered [56,57]. Our interaction analyses focused on surprisal because our hypotheses concerned how cognitive load and TMS alter the processing cost of unexpected input, for which surprisal is a well-established predictor of word-by-word reading times [4,5]. However, entropy was retained to account for context-level predictive uncertainty [58].

### Statistical analysis

#### Reading and reaction times

Mixed effects regression was used to analyse log-transformed reading times and n-back reaction times. We employed LMMs to model parametric terms involving factors such as stimulation condition and task and additionally used GAMMs to explore potential non-linear relationships of continuous predictors such as lexical surprisal and elapsed time across a task with stimulation condition and task.

Separate models were fitted for reading times and n-back reaction times. The reading-time model included all trials from the reading-only and dual-task conditions, whereas the n-back reaction-time model included all trials with a correct n-back button response from the single n-back and dual-task conditions. In both models, fixed effects included stimulation condition (Sham, Offline, Offline+Online TMS), task, and their interaction. Session was included to account for learning effects across visits, and a binary regressor coded whether the respective trial coincided with an online rTMS train. Both models further included participant-wise information including years of education, age, an executive-function composite score, a reading-span score, and session-wise side-effect scores as covariates.

The reading-time model additionally included whether an n-back response in the form of a button press occurred on that trial and whether there was any form of punctuation with the word. Moreover, the following word-specific continuous predictors were included: lexical surprisal, entropy, word frequency, word length, previous-word reading time, and previous-word surprisal. Potential redundancy among psycholinguistic predictors was assessed using variance inflation factors, which were all VIF < 5 for main effects (Table S15). To test whether the effect of lexical surprisal on reading times differed across stimulation conditions and tasks, the reading-time model also included the two-way interactions between lexical surprisal, stimulation condition, and task, as well as their three-way interaction. For the random-effects structure, both reading-time and reaction-time models included a participant-specific random intercept, as well as by-participant random slopes for stimulation condition and task. The reading-time model further included a text-specific random intercept.

GAMMs for reading and reaction times were set up similarly to the LMMs with the goal to test for potential non-linear relationships with continuous predictors such as time across task and lexical surprisal. To this end, elapsed time since task onset and surprisal were included in an interaction with stimulation condition. Interaction terms were specified using identifiable sum-to-zero (sz) smooths together with their required lower-order terms. Moreover, the GAMM allowed to model both surprisal and time effects as by-participant random slopes. Table S2 shows the full equations for LMMs and GAMMs. In the LMMs, if a model formula showed convergence or singularity issues, the random structure was simplified (removing correlations, then removing individual terms) until these warnings disappeared. Following recommended procedures [59], in general the maximally supported random structure was chosen to reduce inflation of Type I errors.

We additionally fitted separate models for reading times and n-back reaction times restricted to trials with an online TMS train. Because these analyses were limited to trials with online stimulation, the offline AG session was excluded. The models were specified analogously to the main models (see Table S2).

Competing LMMs and GAMMs were compared using likelihood-ratio tests for nested models, as implemented in the stats and itsadug packages [60]. All categorical predictors were coded using sum-to-zero contrasts (simple coding). In the GAMMs, continuous predictors were modeled with smooth terms where appropriate, using thin plate regression splines and adequacy of the basis dimension for each smooth was assessed with k.check() from the mgcv package [61]. In the LMMs, all continuous predictors were mean-centered. All models were fit in R v4.6.1 [62] using the packages mgcv [61] and lmerTest [63]. For post-hoc comparisons, the packages performance [64], ggeffects [65], and marginaleffects [66] were used to determine effects between individual factor levels other than the reference levels, with FDR correction applied. Plots were generated using ggplot2 [67], and model output via sjPlot [68].

#### Accuracy in text comprehension questions

Text-comprehension accuracy was analyzed using a binomial GLMM with a logit link, with question-wise responses coded as correct or incorrect. Fixed effects included stimulation condition, task, the interaction of stimulation condition and task, session, block number within session, and the same covariates used in the other models, namely age, years of education, executive-function composite score, reading-span score, and session-wise side-effect score. The random-effects structure included by-participant random intercepts and random slopes for stimulation condition, as well as a random intercept for text. Table S2 displays the model formula. The model was fit using the package glmmTMB [69].

#### Accuracy in n-back trials (d-prime)

D-prime values for the single n-back and dual tasks were analyzed using an LMM, with one d-prime value per task block. The fixed-effects structure was identical to the model for text-comprehension accuracy, including stimulation condition, task, their interaction term, session, block number within session, and the same covariates described above. Because the more complex random-effects structure led to singular fits, the random-effects structure was simplified to include participant-specific random intercepts and random slopes for task, but not for TMS condition. The full model formula is provided in Table S2.

### Computational models for n-back responses

We implemented hierarchical Bayesian DDMs using HSSM v0.4.0 [70] in Python to test whether TMS modulated the latent decision processes underlying n-back performance by jointly modeling reaction times and accuracy. Compared with traditional DDM toolboxes, HSSM supports fully hierarchical model structures analogous to linear mixed models, together with Bayesian likelihood approximation. We specified HSSMs in which the four core DDM parameters, drift rate (*v*), boundary separation (*a*), non-decision time (*t*), and starting point bias (*z*), were estimated simultaneously at both participant and group levels (Fig. 4a).

To account for the potential effect of psycholinguistic predictors on n-back decisions during the dual task, we calculated separate models for single n-back (1- and 2-back) and dual task. Models included trial-wise reaction times above 150 ms and n-back button press choices (correct or incorrect) as dependent variables. The optimal model for each task was determined via model comparison using leave-one-out cross-validation by computing a baseline model, which does not vary any parameter by TMS, task, or participant, and models varying either drift rate alone with TMS, models varying drift rate and either of the other DDM parameters with TMS, and a model varying drift rate, boundary separation, and non-decision time with TMS.

The n-back models included the TMS × task interaction, as well as covariates for session, time within task, and whether the trial received an online TMS train. The random-effects structure included participant-specific random intercepts and random slopes for TMS and task. The dual-task models included the TMS × surprisal interaction, as well as covariates for word entropy, frequency, word length, session, time within task, and a binary indicator of an online TMS train. Participant-specific random slopes for TMS were included. Table S3 provides the full model specifications.

Priors were left at the HSSM default for a safe DDM. Posterior inference was performed using Markov chain Monte Carlo sampling with the No-U-Turn Sampler (NUTS) implemented in NumPyro. Four chains were run in parallel with 1,000 post-warmup draws each, after 2,000 tuning iterations, yielding 4,000 posterior draws in total. Convergence was assessed using R-hat < 1.01 for all parameters, effective sample sizes > 600, and visual inspection of trace plots and posterior density overlays, which are shown in our OSF repository (https://osf.io/79tsa). Factors were sum-to-zero coded, and continuous predictors were mean-centered.

## Data Availability

All behavioral data and the code for the analyses presented in this paper is openly accessible at https://github.com/sandrushba/EXNAT_4_TMS. Derivatives from neuroimaging analyses and computational modeling have been deposited at https://osf.io/79tsa/ and are publicly available.

## Acknowledgments

We thank Alina Leiss, Sophie Thoma, Johannes Kim, Christopher Conrad, and Jim Krups for assistance with recruitment and data collection.

